# Gene deletion studies reveal a critical role for prostamide/prostaglandin F_2α_ synthase in prostamide F_2α_ formation

**DOI:** 10.1101/2020.04.01.020768

**Authors:** Jacques A Bertrand, David F Woodward, Joseph M Sherwood, Alice Spenlehauer, Cristoforo Silvestri, Fabiana Piscitelli, Vincenzo Di Marzo, Maya Yamazaki, Kenji Sakimura, Kikuko Watanabe, Darryl R Overby

## Abstract

Prostamide**/**prostaglandin F synthase (PM/PGFS) is an enzyme with very narrow substrate specificity and is dedicated to the biosynthesis of prostamide and prostaglandin F_2α._ The importance of this enzyme, relative to the aldoketoreductase (AKR) series, in providing functional tissue prostamide F_2α_ levels was determined by creating a line of PM/PGFS gene deleted mice. Deletion of the gene encoding PM/PGFS (fam213b) was accomplished by a two exon disruption. Prostamide F_2α_ levels in WT and PM/PGFS KO mice were determined by LC/MS/MS. Intraocular pressure was measured by tonometry and outflow facility was measured by the *iPerfusion* method in enucleated mouse eyes. Deletion of fam213b had no observed effect on behavior, appetite, or fertility. Intraocular pressure was significantly elevated by approximately 4 mmHg in PM/PGFS KO mice compared to littermate WT mice. No effect on pressure dependent outflow facility occurred, which is consistent with the typically reported prostanoid effect of increasing outflow through the unconventional pathway. The elevation of intraocular pressure caused by deletion of the gene encoding the PM/PGFS enzyme likely results from a diversion of the endoperoxide precursor pathway to provide increased levels of those prostanoids that raise intraocular pressure, namely prostaglandin D_2_ (PGD_2_), thromboxane A_2_ (TxA_2_). It follows that PM/PGFS may serve an important regulatory role in the eye by providing PGF_2α_ and prostamide F_2α_ to constrain the influence of those prostanoids that raise intraocular pressure.

## Introduction

The prostaglandin F_2α_ (PGF_2α_) analogs are established as first line therapies for treating glaucoma. The selective FP receptor agonists were designed by virtue of the need to attenuate the ocular side effects produced by the naturally occurring prostanoid PGF_2α_ (1-3). Similarly, bimatoprost is an analog of prostamide F_2α_ (4, 5). Both PGF_2α_ and prostamide F_2α_ are the products of an identical biosynthetic pathway, but differing with respect to the cyclo-oxygenase substrate. Thus, arachidonic acid is the primary precursor for PGF_2α_, whereas prostamide F_2α_ is biosynthesized from the naturally occurring mammalian endocannabinoid anadamide (6). Following oxidation of arachidonic acid or anandamide, the respective endoperoxide intermediates PGG_2_/PGH_2_ and prostamides G_2_/H_2_, are converted to the active end-products by PGD, PGE, PGF, prostacyclin and thromboxane synthases (7,8). The first enzyme identified and named as a PGF synthase (PGFS) was an aldo-keto reductase (AKR1C3), a broad substrate specificity enzyme (9-13). More recently, a much more substrate specific enzyme and a member of the thioredoxin-like superfamily was discovered and was designated prostamide/prostaglandin F synthase (14,15), PM/PGFS, fam213. This enzyme facilitates a more restricted and targeted formation of PGF_2α_ and prostamide F_2α_ and thereby conferring tighter control of regulatory function(s). Gene deletion is a classical approach for discovering biological functions and the first results of these studies are presented herein, together with the methodology for creating prostamide/prostaglandin F synthase (fam213) knock-out mice. Given the importance of PGF_2α_ and prostamide F_2α_ in the treatment of glaucoma, these present studies focused on regulation of intraocular pressure.

## Methods

### Creation of prostamide/prostaglandin F synthase (fam213) knock-out mice

The prostamide/prostaglandin F synthase (PM/PGFS) knock-out mouse was generated using the RENKA embryonic stem (ES) cell line, derived from C57BL/6N mice (16). To construct a PM/PGFS targeting vector, a 990 bp DNA fragment carrying exons 2 and 3, containing the enzyme active site of PM/PGFS, was amplified by PCR, and inserted into the targeting vector as described previously (17). In this clone, a DNA fragment of a PGK promoter-driven neo-poly (A) cassette was flanked by two Frt sites. These were flanked by two LoxP sites located 139 bp upstream of exon 2 and 298 bp downstream of exon The 5’ side included 5.7 kb of the PM/PGFS gene and the 3’ side 6.5 kb, followed by an MC1 promoter-driven diphtheria toxin (DTA) gene (Fig. 1A). The targeting vector was introduced into RENKA ES cells by electroporation and recombinant clones identified by Southern blot hybridization (Fig. 1B). To produce germline chimeras, recombinant ES clones were microinjected into eight cell stage embryos of CD1 mice. To generate PM/PGFS knock-out mice, chimeric mice were crossed with TLCN-Cre mice, by which recombination was introduced throughout the whole body (18).

**Figure 1.**
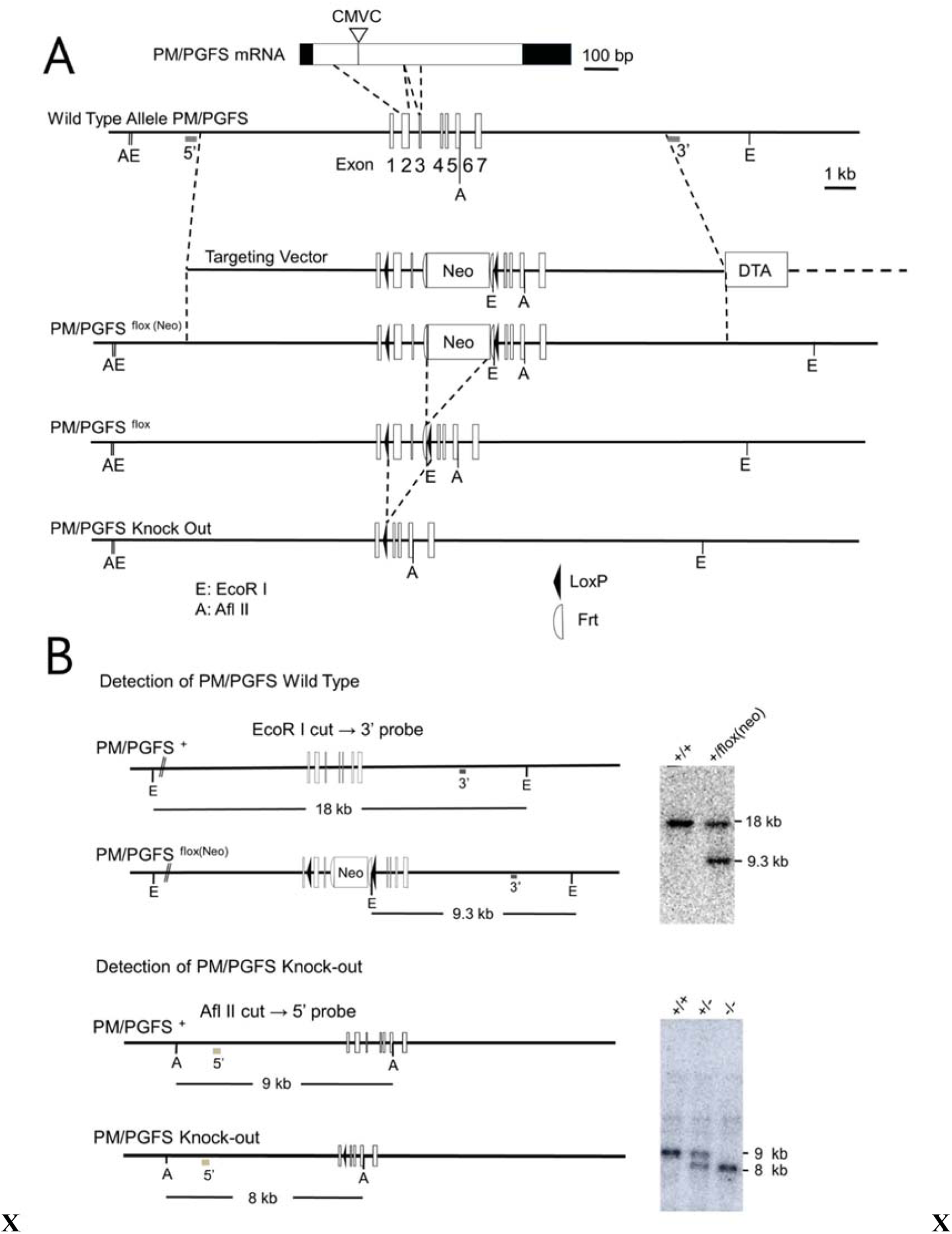
**(A)** A schematic representation of PM/PGFS mRNA, genomic allele, targeting vector, targeted PM/PGFS floxed allele and knock-out allele. Numbered boxes of genomic allele represent the exon sequences of PM/PGFS. Frt sites are indicated by open half-circles. neo, PGK-promoter-neo-pA cassette; DTA, diphtheria toxin cassette; A, Afl II; E, EcoR I. Shadowed boxes indicate 5’- and 3’-Southern blot probes. **(B)** Southern blot analysis of EcoR I- or Afl II-digested genomic DNA prepared from the wild-type (WT), PM/PGFS-floxed (neo) and PM/PGFS knock-out mice.

All procedures on living mice were carried out in compliance with the Imperial College Statement for Use of Animals in Research, under UK home Office project license 70/9064 and in accordance with the ARVO statement for the Use of Animals in Ophthalmic and Vision Reearch. Mice were housed in individually ventilated cages with access to food and water ad libitum, maintained at 21°C with a 12-hour light/dark cycle. Heterozygous mice were used for breeding and all pups genotyped prior to usage. Genotyping was carried out on DNA extracted from ear tissue following manufacturer’s protocols (Express Extract, Kapa Biosystems, Cambridge, MA, USA). TaKaRa Ex Taq polymerase (RR001, Takara Biotech, Shiga, Japan) was used for PCR reactions, forward primer (CAACCAGTCTAACTAGGCT) and reverse primer (CATGTGACCTGAACCCCCT) were cycled 30 times with an annealing temperature of 65°C to yield predicted products of 650 bp for knock-outs and 1500 bp for wild-types. PCR products were resolved by gel electrophoresis (1% agarose) in the presence of a DNA gel stain (SYBR Safe, Invitrogen, Carlsbad, CA). Bands were visualized on an imaging station (Biospectrum 500, UVP, Upland, CA).

### Quantitative real-time PCR

PM/PGFS knock-out, male mice (n=13) and littermate wild type mice of both sexes (n=15), 10 to 15 weeks old, were humanely culled and harvested for eyes and approximately half were also harvested for inguinal, and gonadal white fat pads. Pairs of eyes were cleared of extraocular tissue, hemisected at the equator and lenses removed prior to homogenisation in TRIzol Reagent (ThermoFisher Scientific, Waltham, Massachusetts, USA) with a rotor-stat homogeniser (VDI 12, VWR, Leicestershire, UK). Non-ocular fat pads were placed in TRIzol immediately after dissection, shortly followed by homogenisation. Total RNA was extracted using PureLink RNA spin columns and DNase I treated following the manufacturer’s protocol (ThermoFisher Scientific). RNA yield was determined using a NanoDrop ND-1000 spectrophotometer and Superscript VILO reverse transcriptase (ThermoFisher Scientific) was used to synthesise cDNA, following manufacturer’s instructions. qPCR was carried out on 10 ng of starting RNA using SYBR select master mix (ThermoFisher Scientific). PM/PGFS primers (Table 1) were designed with AlleleID software (Premier Biosoft). cDNA was analysed in triplicate on the QuantStudio 6 Flex (applied Biosystems, ThermoFisher Scientific) Real-Time PCR system. GAPDH was used as the reference gene, RNA content was determined by the delta Ct method of comparative quantification (Ct [reference gene] - Ct [target gene]).

**Table 1.**
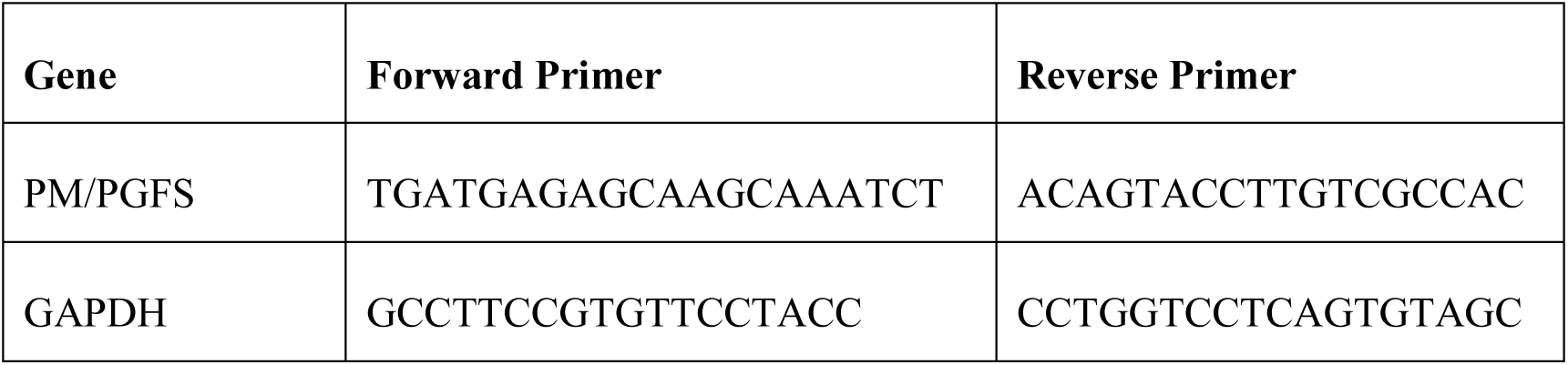
Exon spanning gene expression primer sets for use with SYBR green qPCR. GAPDH was used as a housekeeping gene for relative mRNA abundance.

### Measurement of tissue prostamide F_2α_ levels by LC/MS/MS

PM/PGFS knock-out mice (n=5) and littermate wild type mice (n=5), 10 to 15 weeks old and of both sexes, were humanely culled and harvested for eyes and inguinal and gonadal white fat pads. For prostamideF_2α_, tissues were dounce-homogenized and extracted with acetone containing an internal deuterated standard for prostamideF_2α_ quantification by isotope dilution ([2H]-4 prostamide F_2α_). The lipid-containing organic phase was dried, weighed and pre-purified by open bed chromatography on silica gel. Fractions were obtained by eluting the column with 99:1, 90:10, 70:30 and 50:50(v/v) chloroform/methanol. The 70:30 fraction was used for prostamideF_2α_ quantification by LC-MS-MS, using an LC20AB coupled to a hybrid detector IT-TOF (Shimadzu Corporation, Kyoto, Japan) equipped with an ESI interface. LC analysis was performed in the isocratic mode using a DiscoveryHC18 column (15 cm, 62.1 mm, 5 mm) and methanol/water/acetic acid (53:47:0.05 by vol.) as mobile phase with a flow rate of 0.15 ml/min. Identification of prostamideF_2α_ was carried out using ESI ionization in the positive mode with nebulizing gas flow of 1.5 ml/min and curved desolvation line temperature of 250C.

### Intraocular pressure measurement

PM/PGFS knock-out (n=18) and littermate wild type (n=27) mice, aged 10 to 15 weeks old and of both sexes, received bilateral intraocular pressure measurement between the hours of 10:00 AM and 12:00 noon using a commercial rebound tonometer (TonoLab; Icare, Helsinki, Finland). Intraocular pressure was calculated as the mean of three consecutive tonometer readings as previously described (19). Statistical significance between groups was established using the unpaired Students T-test (mean ± 95% CI). Normal distribution was checked by the Kolgomorov-Smirnov test, equity of variances between groups was determined by Levene’s test.

### Measurement of outflow facility

PM/PGFS knock-out (n=18) and littermate wild type (n=22) mice, 10-15 weeks old and of either sex, were humanely culled by cervical dislocation and eyes enucleated immediately after death. Outflow facility was measured in individual eyes using the *iPerfusion* system, as previously described (20). Briefly, eyes were glued to a support platform submerged in PBS regulated at 35°C. Anterior chambers were cannulated with 33-gauge beveled needles (NanoFil, NF33BV-2, World Precision Instruments) attached to micro-manipulators and equilibrated for 30 minutes at 9 mmHg. The perfusate comprised 0.2 μm filtered DBG (PBS including divalent cations and 5.5 mM glucose). Six pressure steps equally spaced between 6 and 15 mmHg were carried out. For each step, steady state was considered once the slope of the ratio of flow rate to pressure was less than 0.1 nl/min/mmHg/min continuously for 1 minute. Steady state flow *Q* and pressure *P* were extracted by averaging over the last 4 minutes for each step. Pressure steps that failed to reach steady state were excluded from further analysis and eyes with 4 or fewer successful steps were omitted. Outflow facility was estimated by fitting a model of the form

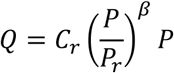

where *C_r_* represents outflow facility at a reference pressure *P_r_* (8 mmHg) and β characterizes the non-linearity of the *Q-P* relationship. Note that there is no term for flow at zero pressure, as this has been demonstrated to be zero (21).

## Results

### Transcription

Transcript for fam213b was undetectable by qPCR in homogenized anterior eye segments and excised inguinal fat pads of homozygous PM/PGFS knock-out mice (Fig. 2). In littermate wild type mice, mRNA was detected in both fat pad and anterior eye segment homogenates.

**Figure 2.**
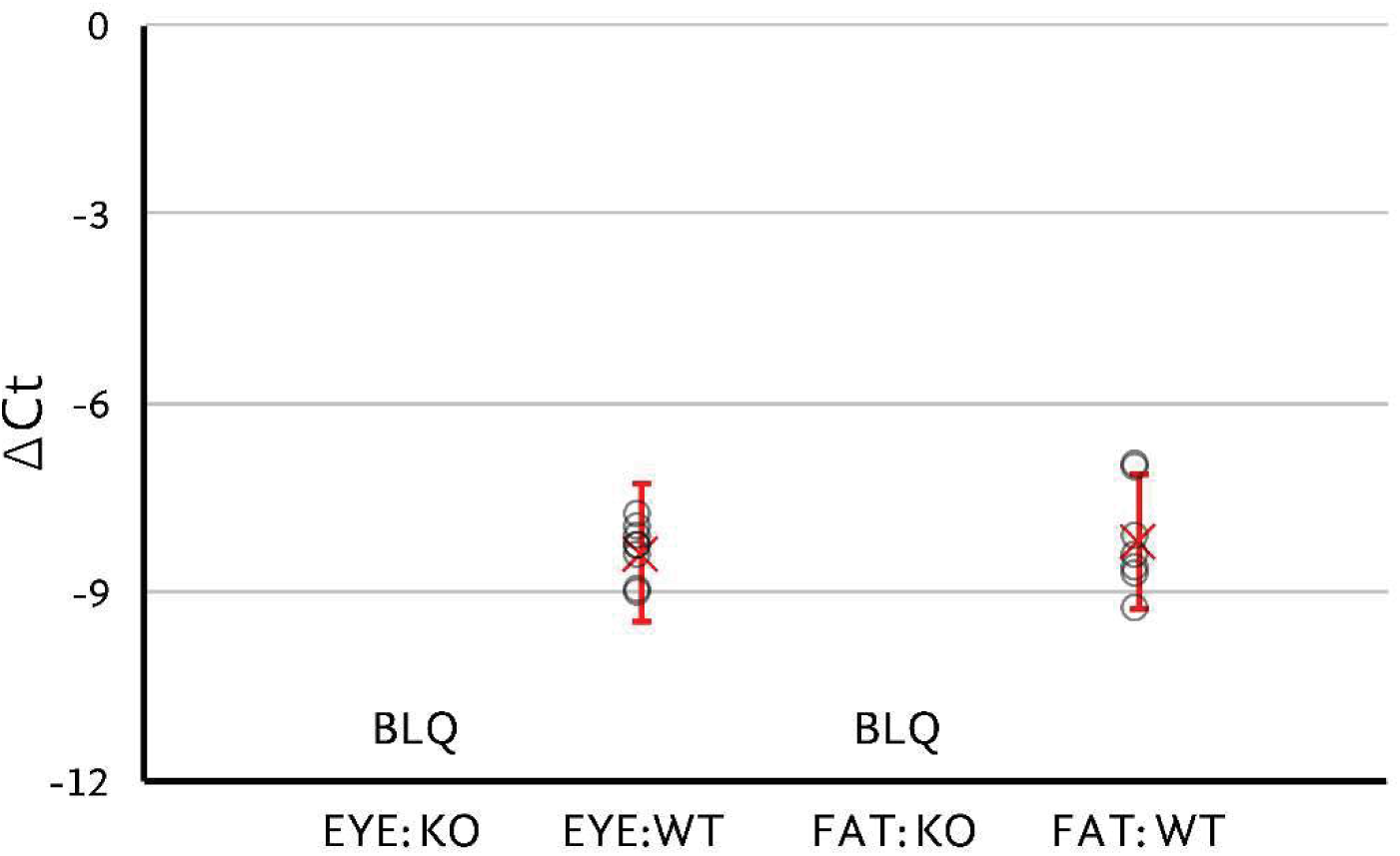
Gene expression analysis of the PM/PGFS by qPCR. PM/PGFS mRNA was undetectable in homogenized anterior segments and inguinal fat pads of PM/PGFS knock-out mice. Similar levels of PM/PGFS mRNA were found in the homogenised anterior segments and inguinal fat pads of littermate wild type mice. Samples were run in triplicate, dCt = Ct [reference gene] - Ct [target gene], with GAPDH as the reference gene, Ct values greater than 35 cycles were excluded. Circles represent individual samples (**○**) and red crosses (X) show sample means with error bars as two standard deviations.

### Tissue prostamide F_2α_ levels by LC/MS/MS detection

Prostamide F_2α_ could not be detected in inguinal fat taken from PM/PGFS knock-out mice, whereas inguinal fat taken from littermate wild type mice yielded low but consistently detectable amounts in all animals (table 2).. In gonadal white adipose tissue and whole eyes, prostamide F_2α_ was never detected in any of the PM/PGFS knock-out or littermate wild type mice. It should be noted that the diminutive mouse eye produces very low yields of ciliary body tissue, which would be further compromised by the immense dilution factor inherent by the presence of additional ocular structures in the anterior segment of the eye.

**Table 2.** Prostamide F_2α_ levels (pmol/g tissue) mean values ± SEM in ocular anterior segment tissues, inguinal white adipose tissue, and gonadal white adipose tissue. No detectable prostamide F_2α_ is designated 0. N=5.

| TISSUE | Wild type mice | Knock out mice |
| --- | --- | --- |
| Ocular anterior segment | 0 | 0 |
| Inguinal white adipose tissue | 1.11 ± 0.56 | 0 |
| Gonadal white adipose tissue | 0 | 0 |

### Ocular effects

Intraocular pressure values obtained from PM/PGFS knock-out and littermate wild type mice are compared in figure 3. Intraocular pressure was significantly elevated by 3.6 [2.9, 4.3] mmHg (*P*<0.0001, N=45, mean [95% confidence interval]) in response to deletion of the PM/PGFS enzyme (Fig. 3A). Outflow facility, as measured in enucleated mouse eyes using the iPerfusion system, was 4.9 [4, 6] nl/min/mmHg (N=18) for PM/PGFS knock-out mice and 5.6 [5, 7] nl/min/mmHg (N=22) for littermate wild-type mice (Figure 3B). The facility at a physiological pressure drop, *C_r_*, was compared between PM/PGFS knock-out and littermate wild type mouse eyes using an unpaired version of the weighted *t*-test of the log-transformed data as described previously (Sherwood *et al*., 2016). Outflow facility was insignificantly different between the two groups (15% [-12, 51%] *P*=0.31, Fig. 3B).

**Figure 3.**
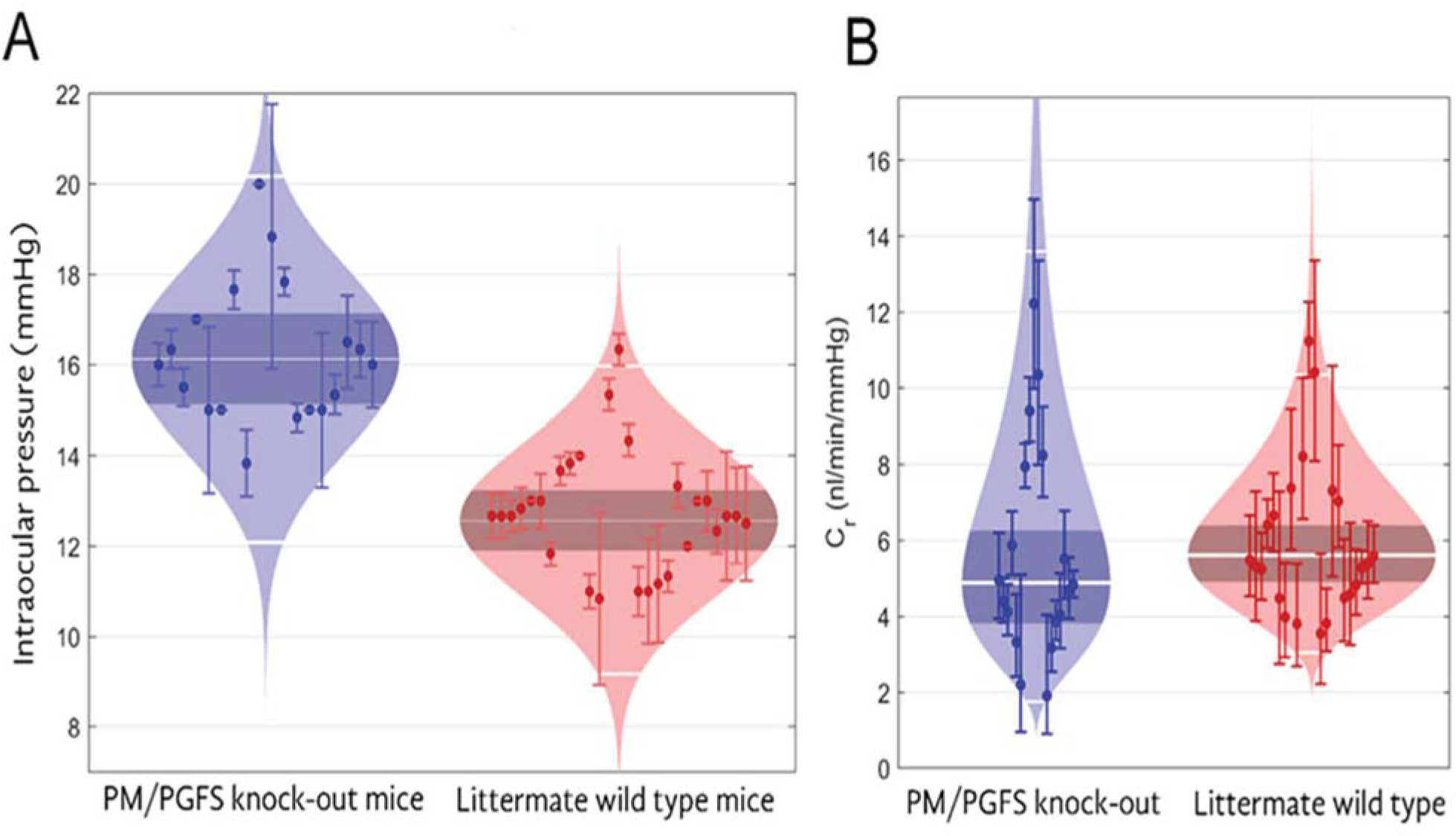
(**A**) Intraocular pressure in PM/PGFS knock-out mice, 16 [15, 17] mmHg (N=18), was significantly greater than littermate wild type mice, 13 [12, 13] mmHg (N=27) (*P*<0.0001). However outflow facility (**B**) was not significantly different, 4.9 [4, 6] vs 5.6 [5, 7] nl/min/mmHg respectively (15% [-12, 51], mean, 95% CI; *P*=0.3, N=22).

## Discussion

Two distinct families of enzymes are known to be involved in the biosynthesis of PGF_2α_ and prostamide F_2α_; the aldoketoreductase family of reductases, notably AKR1C3 (8-13) and PM/PGFS, which is a member of the thioredoxin-like superfamily (14, 15). To date the relative biosynthetic and regulatory importance of these two enzymes in living tissues has remained unclear. By studying prostamide F_2α_ tissue levels in inguinal fat, it became clear that deletion of the gene that encodes PM/PGFS prevents the biosynthesis of prostamide F_2α_. Further, a functional study on intraocular pressure revealed a distinct increase in PM/PGFS knock-out mice, thereby revealing a regulatory function for PM/PGFS for the first time.

PGF_2α_ and prostamide F_2α_ are known to produce profound and long-acting ocular hypotony (1, 22). Despite these pronounced attributes, no evidence has emerged that PGF_2α_ and prostamide F_2α_ are involved in the physiological regulation of intraocular pressure. Deletion of the gene that encodes both the FP receptor and the heterodimeric prostamlde F_2α_ receptor (23, 24) essentially abolishes ocular hypotensive response to prostanoid FP receptor agonist prodrugs and bimatoprost (25). Moreover, in FP receptor knock-out mice, there is no effect on baseline intraocular pressure or the diurnal increase in intraocular pressure that occurs in the early dark phase, 6:00pm – midnight (26). In line with this result in FP knock-out mice is the common knowledge that clinical use of cyclo-oxygenase inhibitors does not result in any meaningful change in intraocular pressure. This is despite the fact that all prostanoids are found endogenously present in the eye (27-29). It seems that under most circumstances prostanoids tending to either increase or decrease intraocular pressure are held in balance and that this balance is not altered, but rather nullified, by global inhibition of prostanoid biosynthesis. It was, therefore, initially considered unexpected that deletion of an enzyme involved in the biosynthesis of PGF_2α_ and prostamide F_2α_ would significantly elevate resting intraocular pressure.

These findings are difficult to explain by the involvement of an aldo-ketoreductase, despite being eminently capable of substituting for absent PM/PGFS by producing PGF_2α_ and prostamide F_2α,_ which are both ocular hypotensive agents. Specifically, one member of the aldo-keto reductase family has been extensively characterized and is designated prostaglandin F Synthase (PGFS) and AKR1C3. In terms of prostanoid formation, it is a dual function enzyme that converts the endoperoxide intermediate PGH_2_ to PGF_2α_ and reduces PGD_2_ to 11β-PGF_2α_ (9, 10), a stereoisomer almost as potent as PGF_2α_ (30). Unlike PGF_2α_, the activity of prostamide F_2α_ is greatly reduced by re-arrangement of the 11-OH group to the β configuration (30). PGFS (AKR1C3) is not expressed in the eye (15). There are, however, alternative enzymes in the aldo-ketoreductase superfamily (31), which could possess PGF_2α_ and prostamide F_2α_ synthase activities. AKR1C1, 2, 3, and 4 share more than 86% sequence identity (32). Nevertheless, the marked increase in intraocular pressure in PM/PGFS knock-out mice, about 20%, suggests that the loss of PM/PGFS is not compensated for by functional expression of AKR1C1-4 in the eye. Although it was not possible to detect prostamide F_2α_ in ocular anterior segment tissue, it was reliably detected from inguinal fat from wild type mice but not from PM/PGFS knockout mice. It appears that prostamide/PGF synthase is the primary source of PGF_2α_ and prostamide F_2α_ since (1) prostamide F_2α_ levels in inguinal fat were not detected in PM/PGFS knock-out mice, and (2) a marked functional effect on intraocular pressure was observed in PM/PGFS knock-out mice.

The absence of PGF_2α_ and prostamide F_2α_ synthesizing enzymes from the ocular anterior segment and a resultant increase in intraocular pressure, would suggest that these prostanoids are involved in its negative regulation. Moreover, the increase in intraocular pressure recorded in PM/PGFS knock-out mice was substantial and similar to that associated with peak levels of diurnal intraocular pressure in mice (26) .This notion would, however, be flawed since disruption of the gene encoding FP and the prostamide F_2α_-sensitive receptors does not alter intraocular pressure (25,26),. A more satisfactory explanation would be the diversion of the endoperoxide precursor PGH_2_ to alternative PG synthases. This would likely be PGD and thromboxane (Tx) synthases, since their products PGD_2_ and thromboxane A_2_ (TxA_2_) have been shown to markedly increase intraocular pressure (33, 34). PGD_2_ has been reported to produce a biphasic effect on intraocular pressure, with an initial ocular marked hypertensive phase (33). TxA_2_ is structurally highly unstable but its selective TP receptor analog U-46619 exclusively produces ocular hypertension that increases over time (34).. No effect on pressure-dependent outflow facility was observed in PM/PGFS knock-out mice, suggesting a possible effect on the uveoscleral pathway, aqueous humor secretion, or both in the ocular hypertension produced by deleting the PM/PGFS gene in mice.

In addition to producing decreased intraocular pressure, an important role for prostamide F_2α_ has been discovered in centrally mediated pain (35) and adipogenesis (36). To date, however, the role of the enzyme PM/PGFS has not been directly studied in regulatory and disease processes. This is the first reported study on PM/PGFS using knock-out mice. The evidence that emerged suggests that this enzyme regulates intraocular pressure not by producing ocular hypotensive PGF_2α_ and prostamide F_2α_ but rather by diverting PGH_2_ conversion to excessive production of PGD_2_ and TxA_2_, both of which cause ocular hypertension. In this series of experiments, the much more substrate specific enzyme PM/PGFS was the important enzyme for prostamide F_2α_ biosynthesis and not AKR1C3.

## Abbreviations

AKR: aldoketoreductase
C: outflow facility
Fa: aqueous flow
Fu: uveoscleral outflow
IOP: intraocular pressur
PG: prostaglandin
PM/PGFS: prostamide/prostaglandin F synthase
PGFS: prostaglandin F synthase
TxA_2_: thromboxane A_2_

## Data Availability

The data described in this manuscript is located at the following institutions. Dept. of Bioengineering, Imperial College London, London, United Kingdom Département de Médecine, Faculté de Médecine, Université Laval, Quebec, Canada Institute of Biomolecular Chemistry, Consiglio Nazionale delle Ricerche, Pozzuoli, Italy Dept. of Cellular Neurobiology, Brain Research Institute, Niigata University, Niigata, Japan Dept. of Nutrition, Koshien University, Takarazuka, Hyogo, Japan.

## Acknowledgements

Y Inoue was a postdoctoral fellow, financially supported by Allergan, who greatly contributed to the creation of a PM/PGFS knock-out mouse. She sadly passed away at a young age after completion of this research.

We gratefully thank Rie Natsume (Brain Research Institute, Niigata University) for her technical assistance. These studies were financially supported by grants from JSPS KAKENHI and Allergan Inc..

